# Morphological and molecular characterisation of *Orientia tsutsugamushi* grown in tick cells

**DOI:** 10.64898/2025.12.15.694307

**Authors:** Magda A. Rogowska-van der Molen, Filomena Gallo, Lesley Bell-Sakyi, Jeanne Salje

## Abstract

*Orientia tsutsugamushi* (*Ot*), the causative agent of scrub typhus, is an obligate intracellular bacterium naturally maintained in *Leptotrombidium* mites, yet its interactions within arthropod hosts remain poorly understood. Here, we developed two tick cell lines, *Ixodes scapularis* ISE6 and *Rhipicephalus microplus* BME/CTVM23, as arthropod models to investigate the intracellular lifecycle of *Ot* strains TA686 and Karp. Both strains efficiently infected and replicated within tick cells, with ISE6 supporting more robust growth. Electron microscopy images revealed that *Ot* maintains its characteristic cytoplasmic, non-vacuolar location in tick cells and exits infected cells by budding off the surface in a membrane-encased structure. Time-course immunofluorescence imaging demonstrated progressive intracellular replication and dynamic expression of *Ot* outer membrane autotransporters ScaA and ScaC, with ScaC enriched early in infection and ScaA at later stages. Metabolic labelling using a clickable methionine analog L-homopropargylglycine showed that high ScaA abundance correlated with reduced translational activity, suggesting a link between ScaA abundance and late-stage or extracellular-like developmental states. The subcellular location of *Ot* in tick cells differs from the characteristic dynein-driven perinuclear clustering observed in mammalian cells, and was not sensitive to disruption of microtubules, indicating a distinct mode of distribution. Together, these findings identify tick cell lines as tractable and biologically relevant arthropod models for studying *Ot* and dissecting host-microbe interactions.

## Introduction

*Orientia tsutsugamushi* (*Ot*) is an obligate intracellular Gram-negative bacterium in the order Rickettsiales found in the salivary glands and ovaries of *Leptotrombidium* mites. Chigger mites are the primary arthropod reservoir and vector of *Ot,* and the bacteria are maintained through mite generations by transovarial transmission. When the infected chigger larvae bite a human, the bacteria can enter through the skin and cause the disease scrub typhus (1–3). This life-threatening condition causes one of the most severe rickettsial diseases in humans, with a median mortality rate of 6% if left untreated (4, 5). Scrub typhus is a leading cause of severe febrile illness in many rural parts of Asia, a region containing two-thirds of the world’s population. Moreover, recent reports of scrub typhus from Latin America, the Middle East and Africa suggest that the disease is distributed more globally than previously recognised (6–8).

The order Rickettsiales comprises two major families – Anaplasmataceae and Rickettsiaceae (9). The former includes *Anaplasma* spp. and *Ehrlichia* spp. that reside within membrane-bound vacuoles inside host cells and are released primarily through exocytosis-like mechanisms or within vesicles, although host cell lysis may also occur (10). The latter includes *Rickettsia spp.* and *Orientia* spp. that replicate directly in the eukaryotic cytoplasm. The intracellular lifecycle of *Ot* has been studied previously in a range of mammalian cell types (11, 12), where it was discovered that *Ot* enters cells through induced clathrin-mediated endocytosis and macropinocytosis and exits the endolysosomal pathway shortly after entry, where it further replicates freely in the host cytoplasm (13). As an obligate intracellular bacterium, *Ot* has lost the ability to synthesise many essential metabolites, such as amino acids, nucleotides and cofactors, making it entirely dependent on host-derived resources to sustain its intracellular growth (14, 15). In mammalian cells, *Ot* traffics to the perinuclear region using dynein-dependent motility and replicates in a microcolony (16). After 3-7 days post-infection, *Ot* traffics to the cell periphery, where it spreads by budding off the surface of infected cells in vesicles in a manner similar to enveloped viruses (17). These extracellular bacteria (EB) are inactive in terms of replication and protein synthesis but can infect subsequent host cells (17, 18). This lifecycle and transmission route are different to the other studied organisms within the order Rickettsiales.

While significant advances have been made in understanding *Ot* biology in mammalian cells, comparatively little is known about its interaction with arthropod host cells. In its natural lifecycle, *Ot* is maintained in *Leptotrombidium* mites, yet the molecular and cellular basis of this persistence remains largely unexplored. The lack of a robust *in vitro* arthropod model has hindered the investigation of bacterial replication and adaptation mechanisms within vector cells. Tick cell lines have previously proven invaluable for studying vector-borne pathogens, helping to define the complex nature of the host-vector-pathogen interactions, including those involving members of the order Rickettsiales, such as *Rickettsia* and *Ehrlichia* species (19–21). Given the close phylogenetic relationships between ticks and trombiculid mites, and the lack of any established mite cell lines, tick cell cultures offer a surrogate model to investigate *Ot* growth in arthropod cells.

In this study, we investigated the ability of *Ot* strains TA686 and Karp to infect and replicate within two tick cell lines, ISE6 (22) and BME/CTVM23 (23). By establishing this system, we aimed to provide a new experimental framework to study the vector stage of *Ot* biology. We compared the two strains in phylogenetically distinct tick cell lines to assess their *in vitro* growth properties and metabolic activity, and characterised the intracellular lifecycle using immunofluorescence microscopy. Altogether, our findings suggest that tick cells represent a promising alternative arthropod model for investigating *Ot* biology.

## Materials and Methods

### Tick cell lines

The *Ixodes scapularis* embryo-derived cell line ISE6 (22) was maintained at 32 °C in L-15B300 medium (24) supplemented with 10% tryptose phosphate buffer (TPB), 10% fetal bovine serum (FBS; Thermo Scientific), 2 mM L-glutamine (Sigma-Aldrich), and 0.1% bovine lipoprotein concentrate (MP Biomedicals). The *Rhipicephalus microplus* embryo-derived cell line BME/CTVM23 (23) was maintained at 32 °C in L-15 (Leibovitz) medium supplemented with 10% TPB, 20% FBS, and 2 mM L-glutamine. Culture media were supplemented with 100 U/mL penicillin and 100 µg/mL streptomycin (Sigma-Aldrich). Both cell lines were grown in 2.2-mL volumes in flat-sided cell culture tubes (Nunc) with weekly medium changes and were split every two weeks as described previously (25).

### Bacterial propagation

*Ot* strains TA686 and Karp were propagated in the mouse fibroblast cell line NCTC clone 929 (L929) purchased from ATCC (Catalogue No. CCL-1) following a previously described protocol (26). Briefly, L929 cells were seeded in T75 flasks at a density of 3 × 10^6^ cells per flask and incubated overnight before infection. A frozen bacterial stock was thawed and added directly to the flasks. On day 6 post-infection, the medium was removed, followed by washing with phosphate buffered saline (PBS; Gibco) once and the cells were then scraped from the flask. The cells were resuspended in plain Dulbecco’s Modified Eagle Medium (DMEM; Thermo Scientific), transferred to a 2 mL Safe-Lock tube and homogenised using a bead-mill homogeniser (Fisher Scientific, Finland) at a speed of 5 for 1 min. The suspension was then centrifuged at 300 × *g* for 5 min. Supernatant was collected, then centrifuged at 14,000 × *g* for 3 min to pellet the *Ot* bacteria. The bacterial pellet was resuspended in sucrose-phosphate-glutamate buffer and stored at −80 °C until use.

### *In vitro* growth in tick cells

Bacteria were added directly to the tick cells in growth media at a multiplicity of infection (MOI) of 50, and incubated for 24 h. After incubation, the supernatant was removed, and fresh growth medium was added to the inoculated cells. Samples for DNA extraction and immunofluorescence microscopy (IFM) were taken by resuspending the cells with a serological pipette. For DNA extraction, 100 µL of cell suspension was added to 900 µL alkaline lysis buffer (25 mM NaOH, 0.2 mM EDTA), and the samples were boiled at 95 °C for 30 min to inactivate bacteria. The samples were stored at −20 °C until further use. For the IFM, µ-Slide 8-well ibiTreat coverslips coated with Concanavalin A (ConA) were prepared before seeding cells as described previously (27). Briefly, the 8-well ibiTreat was coated with 0.5 mg/mL ConA (AA Blocks, USA) working solution for 30 min at room temperature. Then, each well was rinsed with sterile distilled water and dried for at least 16 h before seeding the cells. 5 × 10^4^ cells/mL were seeded in each well of the 8-well ConA ibidi, and incubated at 32 °C for 2 h. For the nocodazole assay, ISE6 cells infected with *O. tsutsugamushi* strain Karp were treated with either DMSO (control) or 10 µM nocodazole dissolved in DMSO (Sigma Aldrich) for 6 h at 32 °C prior to imaging. Then samples were fixed with 4% formaldehyde solution (w/v; Thermo Scientific) for 30 min at room temperature, followed by rinsing three times with PBS. Samples were stored in the dark at 4 °C until staining.

Bacterial concentration was quantified by qPCR (26). The primers and TaqMan probe used for the 47 kDa target gene were as follows: forward 5’-TCCAGAATTAAATGAGAATTTAGGAC-3’, reverse 5’-TTAGTAATTACATCTCCAGGAGCAA-3’, and probe 5’-FAM-TTCCACATTGTGCTGCAGATCCTTC-TAMRA-3’. The qPCR mixture consisted of 1X qPCR Probe Mix LO-ROX (PCR Biosystems, UK), 0.1 µM of each forward and reverse primer, 0.2 µM probe, RT-PCR grade water (Thermo Scientific), and 1 µL of extracted DNA. Real-time PCR was performed on a CFX Duet Thermal Cycler (Bio-Rad, USA) using the following settings: initial denaturation at 95 °C for 2 min, followed by 45 cycles of 95 °C for 15 s, 60 °C for 30 s, with fluorescence acquisition during the 60 °C annealing/extension phase. DNA copy numbers were calculated by comparison with a standard curve constructed from a serial dilution of a 47 kDa standard in the range of 10^1^ to 10^7^ copies (28).

### Scanning electron microscopy (SEM) and transmission electron microscopy (TEM)

*Ot* -infected ISE6 cells at 12 days post-infection and BME/CTVM23 at 19 days post-infection were fixed overnight at 4 °C in a solution containing 2% glutaraldehyde and 2% formaldehyde in 0.05 M sodium cacodylate buffer (pH 7.4) supplemented with 2 mM calcium chloride. Following fixation, samples were washed five times with 0.05 M sodium cacodylate buffer (pH 7.4) and post-fixed in a solution of 1% osmium tetroxide and 1.5% potassium ferricyanide in the same buffer for 3 days at 4 °C. After five washes in deionised water (DIW), samples were incubated with 0.1% (w/v) thiocarbohydrazide in DIW for 20 min at room temperature in the dark. This was followed by five DIW washes and a second osmication step using 2% osmium tetroxide in DIW for 1 h at room temperature. Samples were washed five times again in DIW and block-stained with 2% uranyl acetate in 0.05 M maleate buffer (pH 5.5) for 3 days at 4 °C. After a final series of five DIW washes, samples were dehydrated through a graded ethanol series (50%, 70%, 95%, 100%, 100% dry) and 100% dry acetonitrile, with three exchanges at each step for at least 5 min.

Infiltration was performed overnight in a 1:1 mixture of 100% dry acetonitrile and Quetol resin (without BDMA), followed by 3 days in 100% Quetol (without BDMA). Samples were then infiltrated for 5 days in 100% Quetol resin containing BDMA, with daily resin exchanges. The Quetol resin mixture consisted of 12 g Quetol 651, 15.7 g NSA (nonenyl succinic anhydride), 5.7 g MNA (methyl nadic anhydride), and 0.5 g BDMA (benzyldimethylamine; all reagents from TAAB). Samples were embedded and cured at 60 °C for 2 days.

For SEM imaging, thin sections (∼200 nm) were cut using a Leica Ultracut E ultramicrotome, placed on Melinex coverslips, and air-dried. Coverslips were mounted on aluminium SEM stubs using conductive carbon tabs, and edges were painted with conductive silver paint. Samples were sputter-coated with 30 nm carbon using a Quorum Q150T E carbon coater. Imaging was performed using a Verios 460 scanning electron microscope (FEI/Thermo Fisher) at 4 keV accelerating voltage and 0.2 nA probe current in backscatter mode, using the concentric backscatter detector (CBS) in immersion mode at a working distance of 3.5–4 mm. Stitched maps were acquired using FEI MAPS software with the default stitching profile and 5% image overlap.

For TEM imaging, ultra-thin sections (∼90 nm) were cut and collected on 300 mesh bare copper grids. Samples were imaged using a Tecnai G2 transmission electron microscope (FEI/Thermo Fisher) operated at 200 keV accelerating voltage. A 20 μm objective aperture was used to enhance image contrast. Images were acquired with an AMT digital camera.

### Immunofluorescence microscopy

Cells were permeabilised on ice by incubating in absolute ethanol for 30 min, followed by 0.5% Triton X-100 for 30 min. A blocking step was performed at room temperature in the dark using PBS (pH 7.4) containing 1% bovine serum albumin for 30 min. The samples were then incubated at 4 °C overnight with an in-house-generated primary antibody, i.e. human and rat monoclonal antibodies against TSA56 for strains TA686 and Karp respectively, rabbit monoclonal antibody against autotransporter protein ScaA (#AP84257, Abcam, UK) and rabbit monoclonal antibody against autotransporter ScaC. Β-tubulin was labelled using a mouse primary antibody (T4026, Sigma-Aldrich). The samples were then incubated with a secondary antibody, i.e goat-anti-rabbit Alexa Fluor^TM^ Plus 555 (A32732, Thermo Scientific), goat anti-human Alexa Fluor^TM^ 488, goat-anti-human Alexa Fluor^TM^ 647 (A21445, Thermo Scientific) or goat anti-rat Alexa Fluor^TM^ 488 (A11006, Thermo Scientific) at a dilution of 1:500, supplemented with the nuclear stain DAPI (1:1000; Adva Tech Group Inc. USA) and plasma membrane with HCS CellMask^TM^ Deep Red (1:5000; H32721A, Thermo Scientific) at 37 °C for 1 h. Samples were washed with PBS after each immunofluorescence labelling step. Samples were mounted using mounting medium (20 mM Tris, pH 8.0, 0.5% N-propyl-gallate, 90% glycerol). Imaging was performed on a Nikon Eclipse Ti2-E Inverted Microscope (Nikon, Japan) equipped with a PLAN Apo Lambda 100x Oil (numeric aperture: 1.45) objective plus Intermediate Magnification switching to 1.5X on the main body. 405 nm, 488 nm, 555 nm, and 647 nm filters were applied with a Hamamatsu ORCA-fusion Digital Camera C14440. Z-stack was performed using Define from top to bottom, followed by MaxIP (Maximum Intensity Projection) and 2D Deconvolution.

Click labelling was based on the Click-iT HPG Alexa Fluor protein Synthesis Assay Kit (Thermo Scientific) strategy as described previously (29). Briefly, L-15B300 medium was removed from ISE6 tick cells infected with *Ot* strain TA686 and replaced with minimal DMEM medium lacking L-methionine and FBS and containing 50 µM L-homopropargylglycine (HPG) for 6 h at 32 °C. Bacteria were then washed and fixed with 4% PFA and labelled with primary and secondary antibodies as described above. After the last washing, the Click-iT reaction cocktail was supplemented, and cells were incubated in the dark for 1 h at room temperature. The Azide dye Alexa Fluor 488 was used at a final concentration of 5 µM. The cells were imaged as above.

## Results

### *Orientia tsutsugamushi* replicates in tick cells

Tick cell lines share many physiological characteristics with the arthropods from which they were derived. They can be maintained in mammalian culture media supplemented with mammalian serum, and can grow at incubation temperatures between 28 °C and 34 °C (21). Previous studies have reported the optimal growth temperature for ISE6 and BME/CTVM23 at 32 °C (30), which differs from 37 °C used for most mammalian cell models. Based on this, we compared the growth of *Ot* at different temperatures to determine how temperature influences bacterial replication within tick cells (Fig. 1A). We compared the growth at optimal, i.e. 32 °C, with suboptimal temperatures at 25 °C and 35 °C and determined that there was no difference in *Ot* growth among these temperatures in either ISE6 or BME/CTVM23. However, bacterial replication was more rapid in ISE6 cells compared to BME/CTVM23, although both cell lines supported high bacterial loads. The growth of the avirulent *Ot* strain TA686 was comparable to that of the virulent strain Karp strain at 32 °C (Fig. 1B).

**Figure 1.**
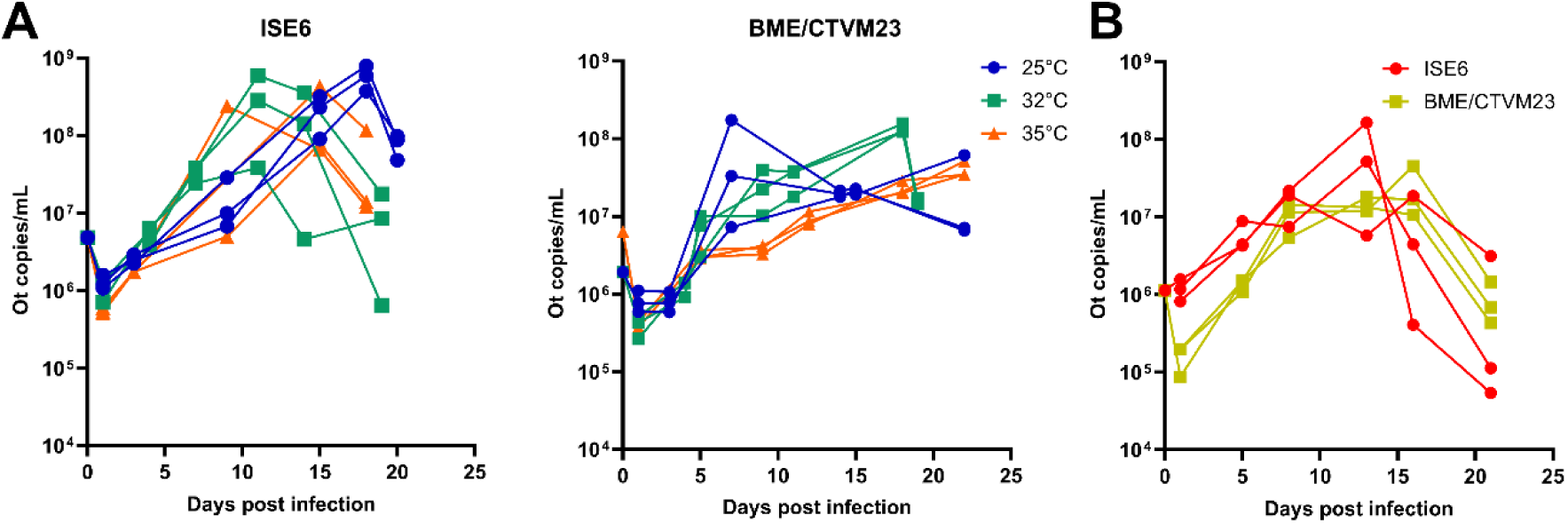
*Orientia tsutsugamushi* grows in tick cell lines. A) Graphs showing bacterial copy number per mL of the *Ot* strain TA686 at 25, 32, and 35 °C in ISE6 and BME/CTVM23 cell lines. B) Bacterial copy number of the *Ot* strain Karp in ISE6 and BME/CTVM23 at 32 °C. Bacterial load was measured by qPCR using primers against the conserved single-copy bacterial gene *tsa47*.

### *Ot* maintains a cytoplasmic location in tick cells

*Ot*, an intracellular pathogen that resides free in the cytoplasm without a host cell-derived vacuole (non-vacuolar), has been previously imaged in invertebrate tissues and mammalian cell lines (31, 32), but no imaging has been reported in arthropod cell lines. Arthropod cells represent a natural environment for vector-borne pathogens; hence, understanding *Ot* intracellular interactions in these cells is essential for unveiling its adaptation and transmission mechanisms. To investigate whether *Ot* localisation and replication in arthropod cells was similar to that described for mammalian cells, we conducted TEM and SEM imaging to characterise its intracellular distribution, morphology and association with host cell structures. TEM and SEM images of *Ot* in ISE6 and BME/CTVM23 cells revealed multiple *Ot* bacteria in the cytoplasm of tick cells that were not enveloped by the host cell membrane (Fig. 2). The bacteria were distributed throughout the host cell cytoplasm with no apparent localisation to specific regions. Two membranes could be observed, as expected for this diderm bacterium, with the inner membrane appearing more electron dense than the outer membrane (Fig. 2). This is in contrast to reports of *Ot* in mite tissue in which electron-dense outer membrane and less-dense inner membrane were seen, although this may reflect differences in sample preparation (32). *Ot* in both tick cell lines measured approximately 0.5 µm in width and 2-4 µm in length. Moreover, in both cell lines, *Ot* appeared to bud off the host cell periphery, similar to its behaviour in mammalian cells (32), acquiring a portion of the host plasma membrane during the process. Some *Ot* also displayed blebs, small vesicles formed by closing cell wall fragments at the cell pole (Fig. 2E), as seen in mammalian cells (33). Although the growth of *Ot* in BME/CTVM23 was somewhat slower than in ISE6, there was no difference in bacterial morphology between the two tick cell lines. These findings demonstrated that *Ot* retains its cytoplasmic location and budding mode of exit in tick cells.

**Figure 2.**
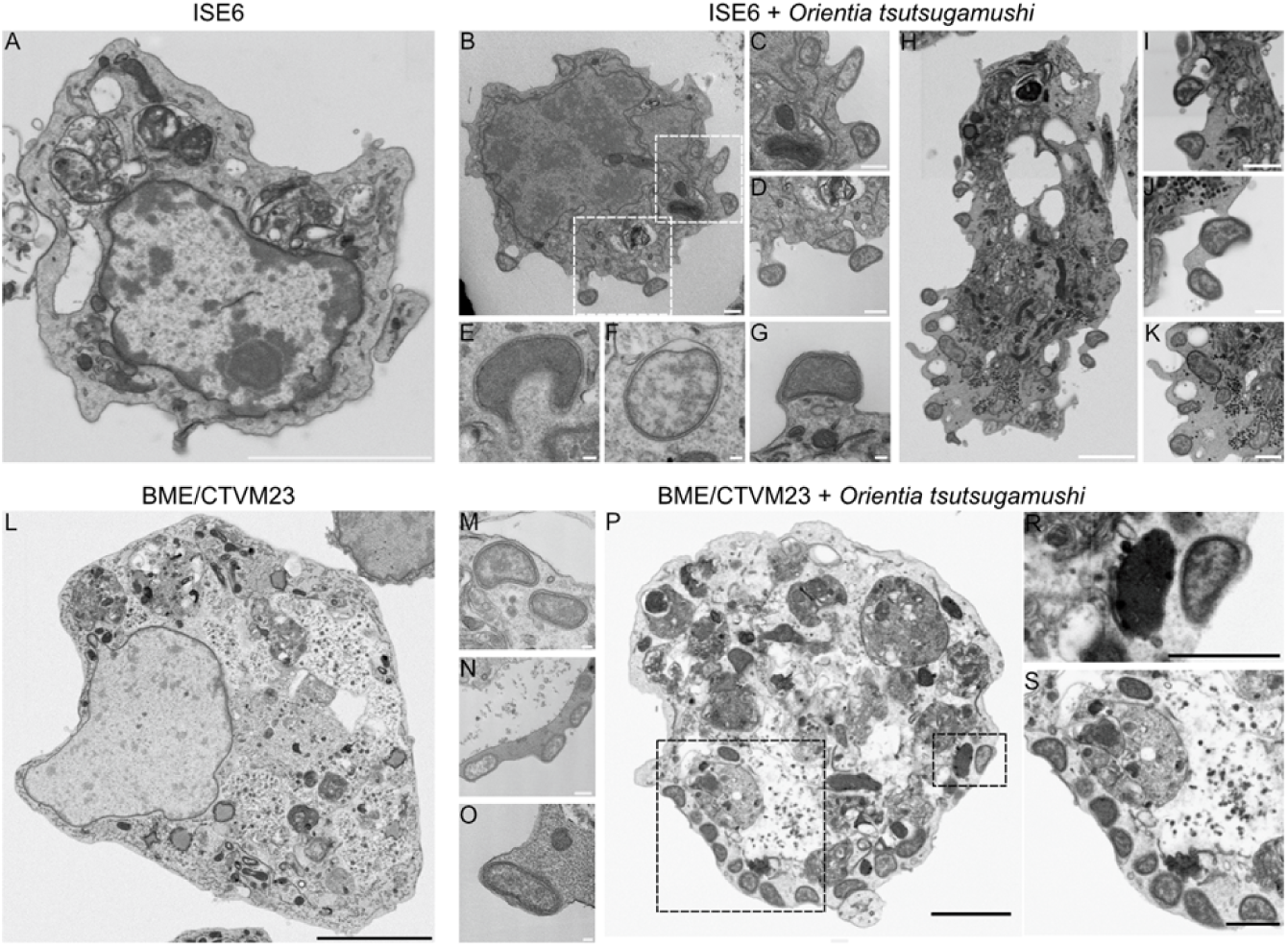
Morphological characterisation of *Orientia tsutsugamushi* grown in tick cell lines. Transmission (A-G, L-O) and scanning (H-K, P-S) electron microscopy images of ISE6 (A-K) and BME/CTVM23 (L-S) tick cell lines, respectively 11 and 19 days post inoculation with *Ot* strain TA686. A) Image of an uninfected ISE6 cell; scale bar 500 nm. B) Image of ISE6 cell infected with *Ot*; scale bar 500 nm. C-D) Detailed images of *Ot* budding from the host cell; scale bar 500 nm. E) Blebbing of an *Ot* cell; scale bar 100 nm. F) Round morphology of *Ot* cell; scale bar 100 nm. G) Budding *Ot*; scale bar 100 nM. H) ISE6 cell infected with *Ot*; scale bar 500 nm. I-K) Budding *Ot*; scale bar 100 nm. L) Image of uninfected BME/CTVM23 cell; scale bar 200 nm. M-O) Detailed images of *Ot* in BME/CTVM23; scale bars 100 nm, 500 nm, 100 nm. P) BME/CTVM23 cell infected with *Ot*; scale bar 500 nm. R-S) Detailed images of *Ot*; scale bar 200 nm.

### The infection cycle of *Ot* in tick cells is characterised by temporal changes in expression of surface proteins

To explore the intracellular growth of *Ot* in tick cells, we performed a time-course infection of ISE6 and BME/CTVM23 cells with strain TA686. Immunofluorescence microscopy revealed that bacterial load increased progressively from 4 to 11 days post-infection in ISE6 and from 5 to 14 days post-infection in BME/CTVM23 (Fig. 3A-D). Whilst most *Ot* strains localise to a tight perinuclear colony in mammalian cells, the avirulent strain TA686 is more dispersed throughout the cytoplasm (34). The subcellular localisation of TA686 in tick cells resembled this previously described dispersed distribution. We used immunofluorescence microscopy to assess expression patterns of three bacterial surface proteins: TSA56, ScaA and ScaC. All three proteins were detected on the surface of at least some bacteria at all time points. The patterns of intensity and distribution of ScaA and ScaC varied at different time points with broadly similar trends between the two cell lines. ScaC expression was observed especially during the early stages of infection (Fig. 3B, D, F), whereas ScaA expression (Fig. 3A, C, E) increased as the infection progressed. ScaA was particularly prevalent on bacterial cells located at the surface of infected cells, especially at late time points. By contrast, TSA56 exhibited relatively stable expression over time, with moderate variation between individual bacteria. On occasion, ScaA and TSA56 displayed heterogeneous expression between individual bacteria within the same host cell, suggesting differential regulation during infection. Likewise, some host cells contained bacteria expressing ScaC but no detectable TSA56 at the early stage of infection (Fig. 3D, F). This temporal difference in expression patterns suggests that ScaC may play a role in early host cell interactions, while ScaA could be associated with a role in maintaining bacterial adhesion or replication. Together, these findings demonstrated that *Ot* TA686 replicates actively within tick cells and dynamically regulates surface protein expression during intracellular development.

**Figure 3.**
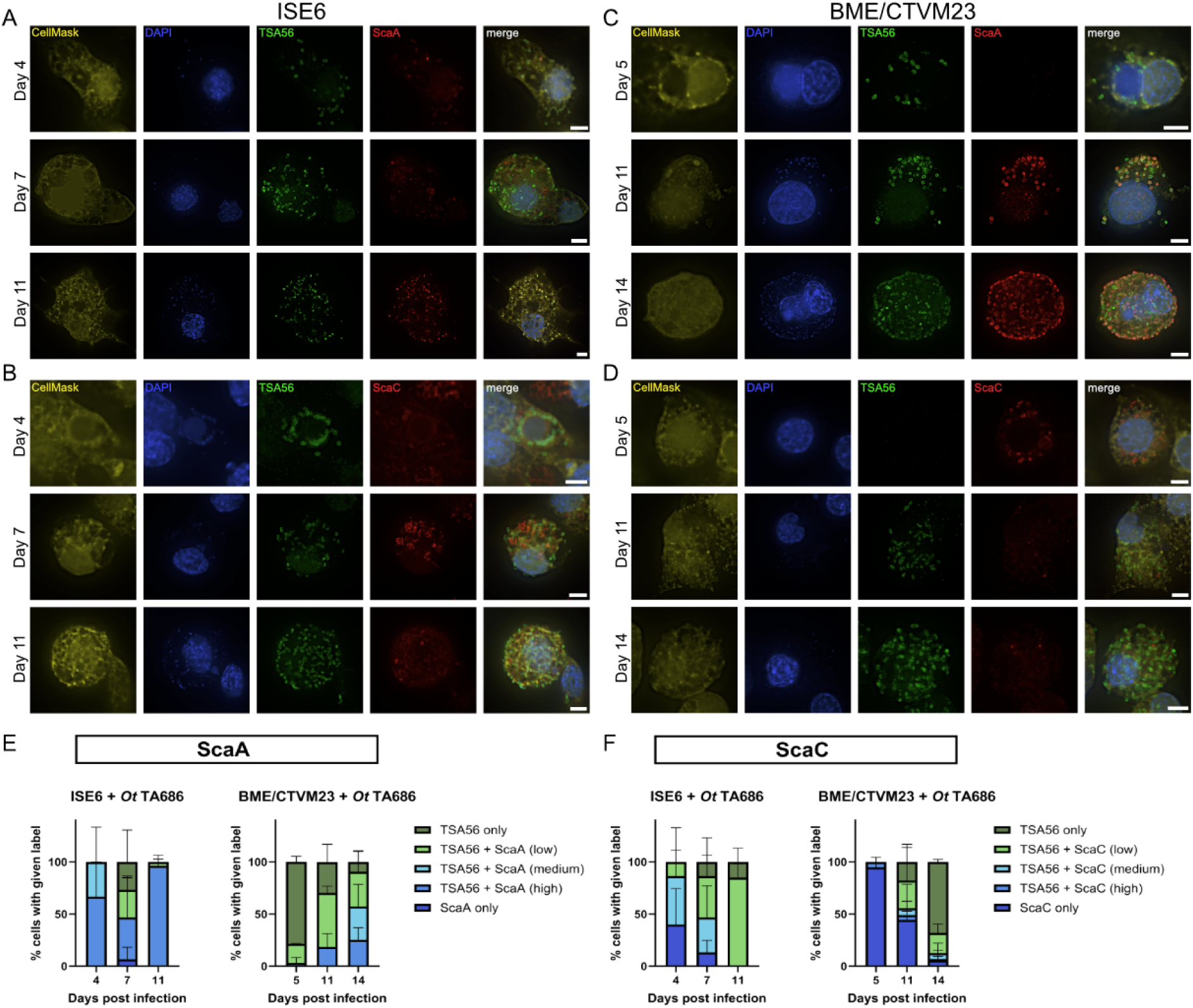
Analysis of the subcellular localisation and expression of bacterial surface proteins TSA56, ScaA and ScaC on *Orientia tsutsugamushi* strain TA686 at different times after infection in ISE6 and BME/CTVM23 cells. A-B) Immunofluorescence microscopy images of *Ot* TA686 in ISE6 at 4, 7 and 11 days post infection. The bacteria expressed the outer membrane proteins TSA56, ScaA (A) and ScaC (B). C-D) Immunofluorescence microscopy images of *Ot* TA686 in BME/CTVM23 cells at 5, 11 and 14 days post-infection. Cytosol and nuclei were stained with CellMask and DAPI, respectively. Scale bars represent 5 µm. E-F) Graphs showing quantification of data represented in A-D. Infected tick cells were scored as having bacteria inside them that were labelled with TSA56, ScaA, ScaC, or both. The data were obtained from three independent biological replicates. A minimum of 20 cells were counted per replicate. Graph shows mean and standard deviation. Low indicates <50%, medium >50% and high 100% of host cells with the given label. All data in this figure were determined using Graphpad Prism software.

### Karp replicates in tick cells and shows a temporal shift from ScaC to ScaA expression

*Ot* strains differ in pathogenicity (34); therefore we sought to analyse the intracellular lifecycle of the virulent Karp strain in ISE6 and BME/CTVM23 cells. Similarly to the avirulent TA686 strain, *Ot* Karp readily infected and replicated within tick cells, with consistent increase in the abundance of TSA56 between day 5 and day 9, indicating active intracellular replication of Karp in both cell lines (Fig. 1B). We explored the subcellular localisation of Karp, and expression of TSA56, ScaA and ScaC proteins (Fig. 4). We found that Karp displayed patterns of ScaA and ScaC expression similar to those seen for TA686, with ScaC decreasing over time and ScaA increasing. The shared pattern of Sca genes expression between TA686 and Karp strains indicates that phase-specific regulation is conserved across phylogenetically distinct strains despite differences in pathogenicity. Furthermore, unlike reports from various mammalian cell lines, we did not observe the characteristic perinuclear colony formation. Instead, the intracellular distribution of the Karp strain resembled that of TA686, with even dispersal throughout the host cell cytoplasm.

**Figure 4.**
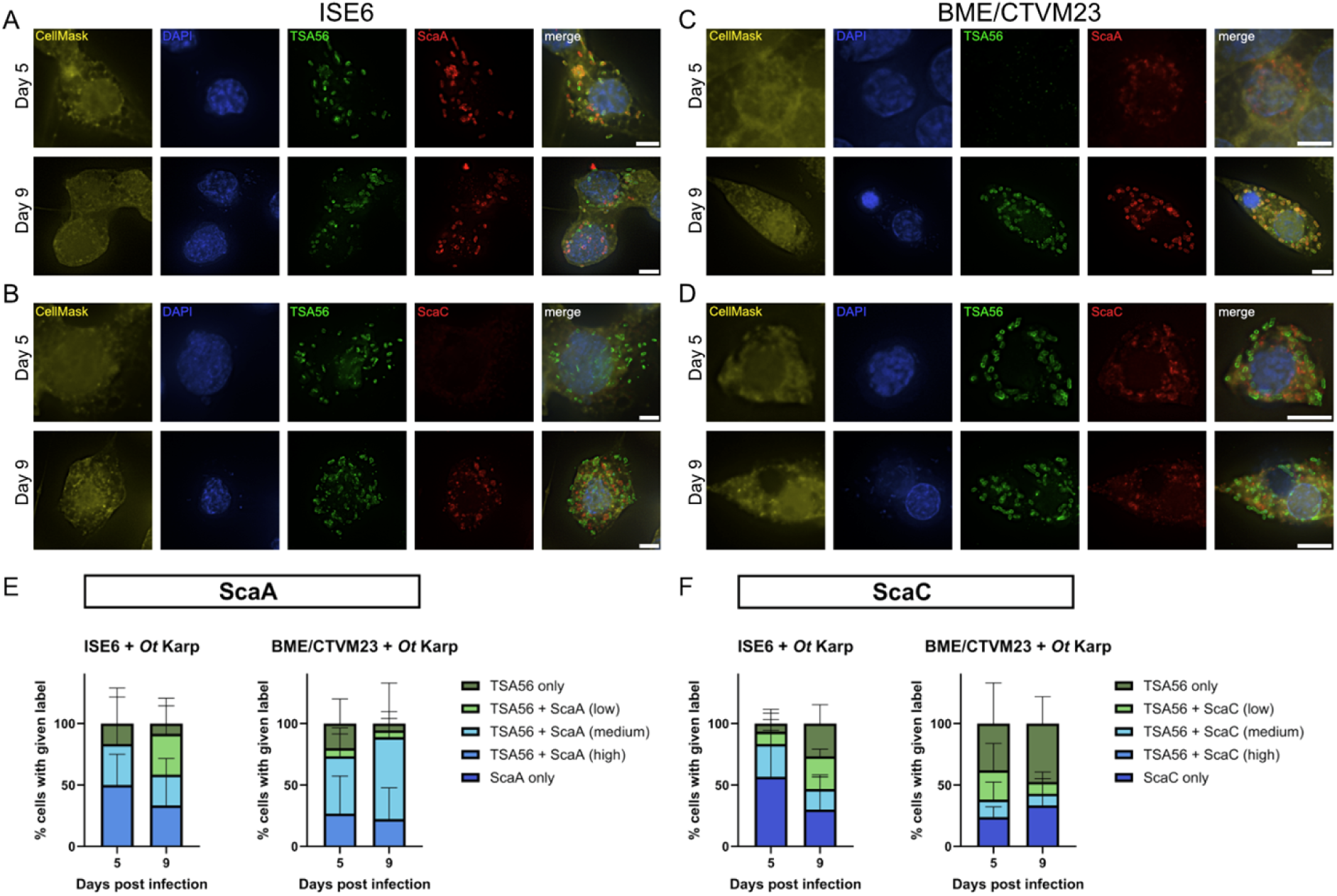
Analysis of the subcellular localisation and expression of bacterial surface proteins TSA56, ScaA and ScaC on *Orientia tsutsugamushi* strain Karp at different times after infection in ISE6 and BME/CTVM23 cells. A-D) Immunofluorescence microscopy images of *Ot* Karp in ISE6 (A-B) and BME/CTVM23 (C-D) at 5 and 9 days post infection. The bacteria expressed the outer membrane proteins TSA56 (green), ScaA (red); A, C), and ScaC (red; B, D). Cytosol (yellow) and nuclei (blue) were stained with CellMask and DAPI, respectively. Scale bars represent 5 µm, *N* = 3, *n* = 20. E-F) Graphs showing quantification of data represented in A-D. Infected tick cells were scored as having bacteria inside them that were labelled with TSA56, ScaA, ScaC, or both. The data were obtained from three independent biological replicates. A minimum of 20 cells were counted per replicate. Graph shows mean and standard deviation. Low indicates <50%, medium >50% and high 100% of host cells with the given label. All data in this figure were determined using Graphpad Prism software.

### Microtubule disruption does not alter *Ot* localization in tick cells

*Ot* typically clusters into a tight perinuclear colony in infected mammalian cells, although it was recently shown that the avirulent strain TA686 is unusual in displaying this phenotype more weakly than other strains (34). It is known that the perinuclear trafficking is mediated by the bacterial surface protein ScaC, that interacts with dynein via a BicD1/2 adapter to move along microtubule filaments (16). Nocodazole treatment, that disrupts microtubule filaments, has been shown to displace intracellular *Ot* from the perinuclear location in human and murine cells *in vitro* (16, 35). The observed lack of perinuclear localisation of either Karp or TA686 *Ot* in the present study led us to ask whether microtubules play any role in positioning of *Ot* in arthropod cells. To address this, we treated *Ot-*infected ISE6 cells with nocodazole and assessed the impact on intracellular bacterial distribution. (Fig. 5). Whilst nocodazole clearly destabilises and depolymerises β-tubulin, a key component of microtubules, in ISE6 tick cells, as shown by a loss of ß-tubulin labelling, there was no change in spatial arrangement of *Ot* Karp. This indicates that *Ot* distribution in tick cells does not require intact microtubules.

**Figure 5.**
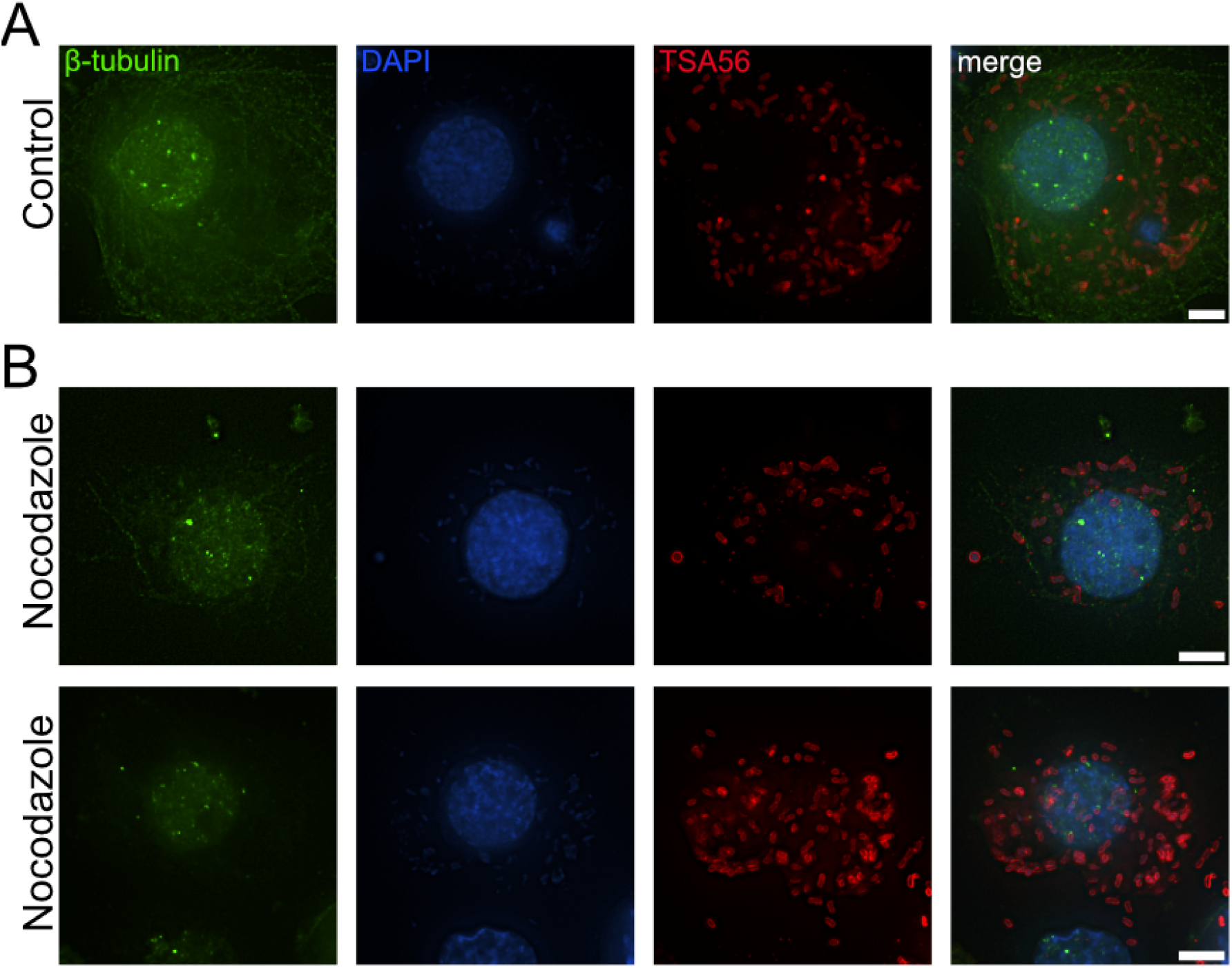
Nocodazole treatment does not alter the intracellular position of *Orientia tsutsugamushi* strain Karp in ISE6 cells. Immunofluorescence images of ISE6 cells at 11 days post-infection, treated with either (A) DMSO (control) or (B) 10 µM nocodazole. Karp was detected using an antibody against the outer membrane protein TSA56 (red); β-tubulin was stained with a primary antibody (green), and nuclei with DAPI (blue). Scale bars represent 5 µm.

### ScaA expression correlates with decreased metabolic activity

*Ot* is a strictly obligate intracellular bacterium with limited metabolic and biosynthetic capability, relying on host cells as sources of carbon, nitrogen and essential metabolic intermediates (36). *Ot* exhibits distinct intracellular (IB) and extracellular (EB) bacterial populations and previous studies showed that, unlike inactive EB, IB are translationally active at five days post-infection in mammalian cells (10). Our observations revealed an increased expression of the ScaA gene during the late stages of infection in tick cells, leading us to hypothesise that ScaA-enriched bacteria are entering a maturation stage of growth in preparation for exit via budding. Since extracellular bacteria are metabolically inactive, we would expect these surface-localised, ScaA-positive bacteria to have decreased metabolic activity. To test this, we used a microscopy-based assay of protein synthesis activity using a clickable methionine analog L-HPG which is conjugated to a fluorophore following fixation (29).

The distribution of bacteria exhibiting different combinations of ScaA abundance and HPG incorporation at 11 days post infection in ISE6 cells is shown in Figure 6. As observed previously, TSA56 and ScaA were heterologously expressed, with substantial variability in translation activity marked with HPG labelling. *Ot* displayed multiple phenotypic states, all expressing TSA56 but with ScaA levels ranging from low to high. Notably, a large proportion of the bacteria lacked ScaA expression (Fig. 6B), and simultaneously, these bacteria showed the highest metabolic activity. In contrast, bacteria with high ScaA abundance were metabolically inactive, exhibiting little to no HPG incorporation. Among the double-labelled populations, most ScaA-positive bacteria displayed only low to medium HPG incorporation, whereas bacteria with both high ScaA and high HPG signal represented only a minor fraction. Together, these findings indicate that high ScaA abundance is associated with reduced protein biosynthesis, supporting the idea that ScaA-enriched bacteria are in a metabolically inactive state characteristic of late-stage developmental transitions prior to cell exit.

**Figure 6.**
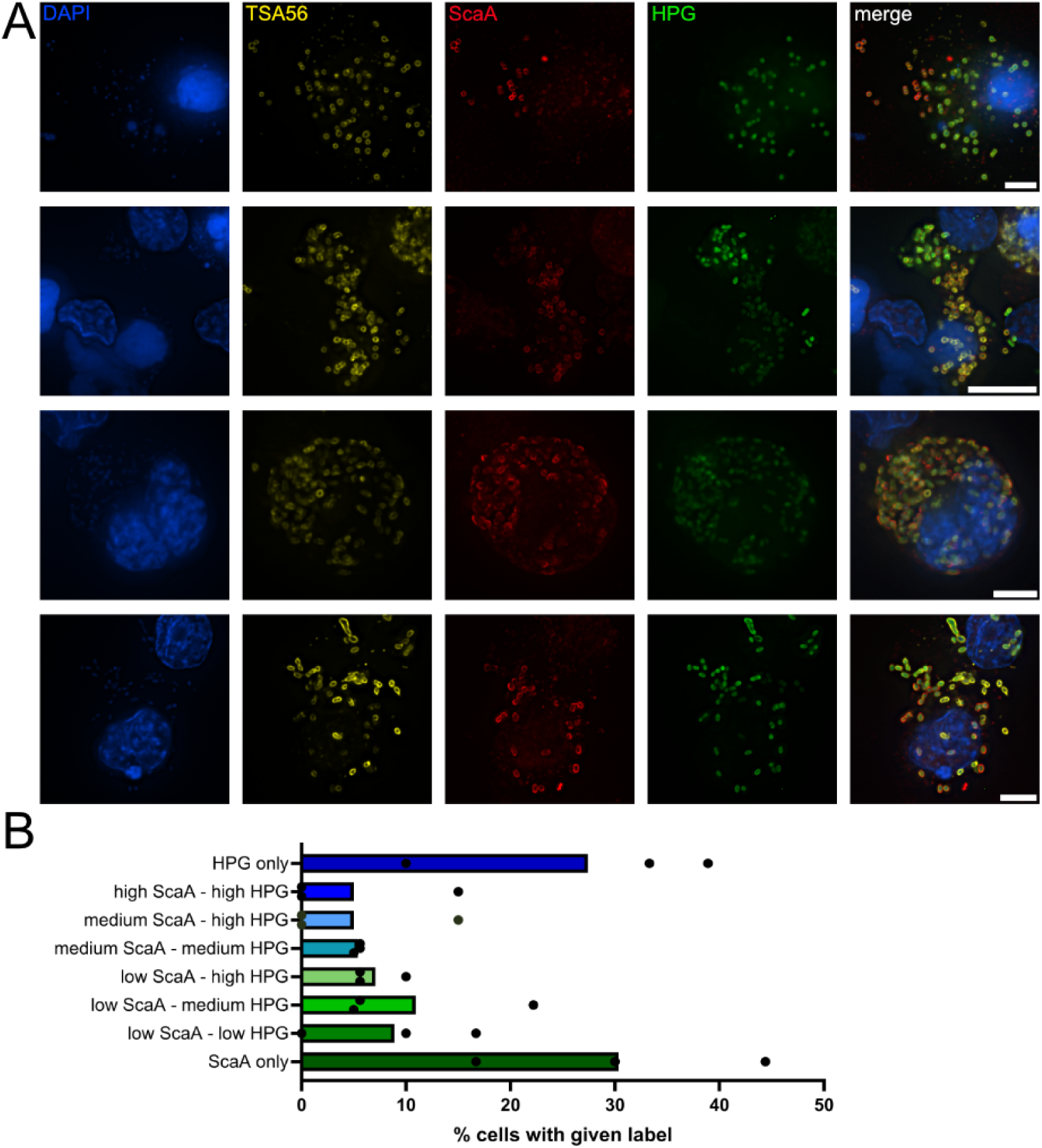
Analysis of the metabolic activity of individual intracellular *Orientia tsutsugamushi* bacteria reveals a negative correlation between ScaA expression and metabolic activity. A) Immunofluorescence microscopy images of ISE6 cells infected with *Ot* TA686 at 11 days post-infection and labelled with the clickable methionine analog homopropargylglycine (HPG, green), antibodies against surface protein TSA56 (yellow) and ScaA (red). DNA (DAPI) is shown in blue. Scale bars represent 5 µm. B) Quantification of metabolic activity, as measured by a detectable HPG incorporation, is shown as the percentage of *Ot*-infected cells falling into each category of combined ScaA abundance and HPG signal. The data were obtained from three independent biological replicates. A minimum of 20 cells were counted per replicate. All data in this figure were determined using Graphpad Prism software.

## Discussion

Under natural conditions, *Ot* is maintained in the *Leptotrombidium* mite population through transovarial transmission, with infection passing directly from females to their offspring. The chigger larvae thus emerge already carrying *Ot*, serving as the primary vectors responsible for transmitting the pathogen to vertebrate hosts. Although *Ot* has been studied *in vivo* in infected mites and mice and *in vitro* in mammalian cell cultures (10, 32, 34), it has not been investigated in any arthropod cell culture system. In this context, our study has unveiled a novel arthropod model for studying this obligate intracellular bacterium, *Ot*, providing new opportunities to dissect vector-pathogen interactions in a controlled and tractable system.

Tick cell lines have an invaluable role in *in vitro* studies to investigate Rickettsiales bacteria. Numerous studies have demonstrated that several vector-borne pathogens can be successfully isolated, cultured and propagated in tick cells, including bacterial species *Anaplasma phagocytophilum*, *Anaplasma marginale*, *Ehrlichia canis*, *Ehrlichia ruminantium*, *Rickettsia raoultii* and a close relative of *Ot*, *Occidentia massiliensis* (21, 23, 30, 37–40). These systems have provided essential insights into pathogen replication dynamics, host cell responses, and mechanisms of vector competence. Likewise, they have also led to comparative studies across different Rickettsiales species, enabling the identification of intricate strategies that these obligate intracellular bacteria employ to survive within arthropod hosts (41, 42). Although *Ot* is a mite-borne bacterium, there are currently no mite-derived cell lines available. This limitation has created a major gap in our ability to dissect primary *Orientia-*host interactions. In the absence of a homologous arthropod model, tick cell lines are an important tool for studying *Ot in vitro*. Tick cells share key innate immune pathways, endocytic processes, and intracellular trafficking machinery with other arthropods, making them valuable for studying how *Ot* invades, persists, and manipulates arthropod cells. In this study, we used two cell lines derived from different tick genera, *I. scapularis* ISE6 and *R. microplus* BME/CTVM23, to investigate *Ot* infection dynamics in arthropod cells.

The observed tick cell infection phenotype differed from that described in mammalian cells. *Ot* exhibited slower growth in both tick cell lines, which is consistent with expected differences in optimal replication temperature between arthropod and mammalian hosts. Interestingly, however, *Ot* growth was not affected by the range of temperatures we tested, namely 25 – 35 °C. This observation suggests that *Ot* may possess a broad thermal tolerance or dynamic adaptive mechanisms that modulate replication independently of temperature variations. This is supported by a previous study on the *Ot* strain Okazaki, which exhibited better growth at 31 °C than 36 °C (43) in mammalian cells, indicating that some *Ot* strains may be intrinsically adapted to cooler conditions. In nature, *Ot* and many other Rickettsiales reside in ectothermic mites and ticks, where environmental temperature can fluctuate considerably. Such flexibility may represent an evolutionary adaptation that enables *Ot* to maintain stable infections within arthropod hosts despite highly variable thermal environments. Furthermore, strains TA686 and Karp showed similar overall infection dynamics in both tick cell lines, but TA686 consistently reached higher numbers than Karp in ISE6 cells. These differences indicate that TA686 replicates more efficiently in tick cells than Karp, suggesting strain-specific variation in adaptation to arthropod hosts.

While *Ot* exhibits apparent differences in its growth dynamics in tick cells, its overall morphology, location free in the cytoplasm and budding resemble the characteristics in mammalian cells (10, 44, 45). This similarity indicated that *Ot* maintains a stable intracellular lifestyle despite the physiological differences between the arthropod and mammalian hosts, highlighting its broad host adaptability.

Immunofluorescence microscopy using three bacterial surface proteins, and microscopy of metabolically active cells using HPG labelling, revealed heterogeneity in *Ot* populations over time and within individual infected cells. In tick cells, *Ot* showed high ScaC expression in early infections, whereas ScaA became predominantly expressed at later stages. It was recently shown that the bacterial autotransporter protein ScaC recruits dynein via adaptor proteins BICD1/2, driving microtubule-based movement (16). Here we showed that *Ot* is localised throughout the cytoplasm rather than in a perinuclear cluster, and that disrupting microtubules with nocodazole does not noticeably alter *Ot* localisation in tick cells. This suggests that ScaC-mediated trafficking may not be used in these cell lines. The increase in ScaA at late times after infection and in bacteria located at the surface of cells suggest a possible role for this protein in maturation and exit. This is supported by the observation that ScaA-positive bacteria at the surface of infected cells generally exhibited decreased translational activity as measured by HPG incorporation, although the substantial heterogeneity in these measurements demonstrates that additional levels of regulation are likely involved.

In summary, by establishing an arthropod cellular infection model for *Ot*, our study provides a critical tool for understanding how this obligate intracellular pathogen interacts with its vector. Insights into the intracellular dynamics, host-specific gene regulation, and maturation processes of *Ot* can inform on strategies to disrupt transmission from mites to humans, ultimately contributing to better prevention and control of scrub typhus. Moreover, understanding vector-borne pathogen biology in arthropod cells can guide the development of vaccines, addressing a significant global health threat.

## Acknowledgements

We thank the University of Minnesota for permission to use the ISE6 cell line and the Tick Cell Biobank at the University of Liverpool, and Dr Catherine Hartley for supplying both tick cell lines and training in their use. We thank the Electron Microscopy Facility of the Microscopy Bioscience Platform for their technical support and training on equipment use. JS was supported by Wellcome Trust grant 224277/Z/21/Z. LBS was supported by the Wellcome Trust grant no. 223743/Z/21/Z.

## Notes

### Competing Interest Statement

The authors have declared no competing interest.

